# Infection dynamics and virulence potential of clinical *Pseudomonas aeruginosa* isolates in human airway epithelial models

**DOI:** 10.1101/2025.04.11.644308

**Authors:** Claudia Antonella Colque, Signe Lolle, Ruggero La Rosa, Maria Pals Bendixen, Asha M. Rudjord-Levann, Filipa Bica Simões, Kasper Aanæs, Søren Molin, Helle Krogh Johansen

## Abstract

Persistent bacterial infections constitute an increasing health problem, often associated with antibiotic resistance. However, despite extensive antibiotic treatment, *Pseudomonas aeruginosa* persists for decades in people with cystic fibrosis (pwCF), remaining susceptible. Host-pathogen interactions during infection may, therefore, be a major contributing factor to treatment failure. Using an infection model based on human airway epithelial cells, cultured at the air-liquid interface (ALI), we simulated the infection process of *P. aeruginosa* to investigate the colonization dynamics and virulence potential during infection in ALI models from non-CF and CF donors, and the BCi-NS1.1 cell line. Infections by reference strains and clinical isolates from pwCF revealed four infection clusters based on virulence, epithelial damage, and localization within the epithelium, in a strain-specific manner regardless of the type of ALI model or clonal lineage. Modulator treatment to restore CFTR channel function did not change infection patterns in CF ALI models. Dual RNA-seq revealed that bacterial colonization of ALI models significantly upregulated host inflammatory pathways, dependent on the strain’s virulence. Simultaneously, while bacterial gene expression was similar in non-CF and BCi-NS1.1 ALI models, CF models promoted differential regulation in a type III secretion mutant. Altogether, our results profile key infection dynamics occurring in pwCF and provide ground knowledge on the interplay between the airway epithelium and *P. aeruginosa*.

## Introduction

With increasing lifespans and changes in lifestyle, persistent bacterial infections are on the rise (*1*). Although antibiotics are generally effective, diagnosing and treating these infections is often challenging because of their complexity and the continuous changes in both the host and pathogen (*2*). This is worsened by the lack of robust and reliable model systems to predict infection outcomes and treatment efficacy. However, by understanding treatment failures in physiologically relevant model systems, improved diagnostics and therapies can be developed (*3*).

People with cystic fibrosis (pwCF) often suffer from persistent bacterial infections caused by a plethora of bacterial pathogens that colonize the airways and persist for decades. Cystic fibrosis (CF) is caused by mutations in the cystic fibrosis transmembrane conductance regulator protein (CFTR), which is crucial for fluid homeostasis (*4*). CFTR mutations result in impaired chloride secretion, increased mucus viscosity and reduced cilia beating, ultimately leading to reduced mucociliary clearance (*5*). This creates the perfect environment for pathogenic bacteria, such as *Pseudomonas aeruginosa*, which is the leading cause of respiratory morbidity and mortality in pwCF (*6*, *7*). Recent advances in CFTR modulator therapies, particularly elexacaftor/tezacaftor/ivacaftor (ETI), significantly improve the well-being in pwCF by correcting and potentiating the function of the CFTR channels (*8*). While it has been shown that during ETI treatment bacterial pathogen counts decrease (*9*), *P. aeruginosa* often persists indicating the need for a deeper understanding of the relationship between ETI treatment and *P. aeruginosa* infections in pwCF (*10–12*).

Early infections are caused by environmental *P. aeruginosa* strains which adapt, over the years, to the CF host environment through genetic, metabolic, and phenotypic changes (*13*, *14*). This enhances antibiotic resistance, immune escape, a biofilm lifestyle and reduces virulence factor production (*15–18*). In most patients, the initial clone drives within-host adaptation. Person-to-person transmission has also been documented in pwCF, and especially *P. aeruginosa* clones such as the DK01 and DK02 (Denmark), and high-frequent clones such as the C clone (ST17 clone, present in Europe and North America and persistent in pwCF) and PA14 (ST253 clone, found worldwide with affinity to CF and non-CF infections) have this capability (*19–25*). However, how colonization and infection dynamics evolve during infection from a *P. aeruginosa*-human airway point of view remains largely unclear. This includes 1) the role of virulence factors, toxins, and specific dominant strains in tissue invasion and immune escape, 2) the epithelial response and long-term effects on epithelial integrity and function, and 3) how modulator treatment changes the infection pattern and the interaction between the airway epithelium and the pathogen. Understanding the early stages of colonization, therefore, may shed light on the bacterial pathogenic potential, antibiotic treatment failure, and allow exploration of new treatment options. In addition, lack of reliable *in vitro* models which recreate the microenvironment for studying infection processes in the airway limits our understanding of colonization dynamics and bacterial virulence potential within human tissues (*26*).

Host-pathogen interactions in the human airways have traditionally been investigated in monolayer cell cultures, most of which are based on cancer cell lines, primary cells or tissue explants (*27*). These present high genomic instability, challenges for their isolation and maintenance, and lack the spatial context required to accurately mimic the cellular and molecular markup of human airway epithelium. To overcome these limitations, we employed 3D air-liquid interface (ALI) models based on the BCi-NS1.1 cell line and primary cells from CF and non-CF donors, which closely mimic the human airway epithelium, exhibiting differentiation, cell polarization, mucus production, and ciliary beating, which are fundamental parameters in understanding infection dynamics and colonization processes (*28*, *29*).

Recent studies have used ALI systems to shed light on *P. aeruginosa* infection processes, resistance mechanisms, and metabolic adaptation (*30–37*). Specifically, infection dynamics and localization of the bacteria within the epithelium can influence both host response and antibiotic susceptibility/resistance (*38*). Similarly, bacterial metabolic processes can influence the host immune response (*17*). However, these studies used laboratory strains which do not reflect the complexity of clinical CF isolates and their infection dynamics. Therefore, modeling infection patterns in physiologically relevant models with clinically relevant strains is of high priority to understand treatment failure and overcome bacterial persistence (*3*).

In this study, we used ALI models to profile *P. aeruginosa* infection using 12 clinical strains isolated from pwCF from frequent and distinct lineages and compared them with well-characterized reference strains. We determined the overall host response to the bacterial infections, including epithelial integrity, cytotoxicity, cytokine secretion, and bacterial colonization patterns in *P. aeruginosa*-infected ALI models generated with non-CF or CF primary cells, or the human telomerase reverse transcriptase (hTERT) immortalized airway basal cell line BCi-NS1.1. Additionally, we determined the impact of the CFTR modulator ETI on *P. aeruginosa* infection pattern. Dual RNA-seq analysis unfolded host-pathogen interactions at the molecular level. Our results highlight the importance of host-pathogen model systems to understand infection dynamics and promote the development of new therapies.

## Results

### Host response upon bacterial infections in ALI models

To study the early dynamics of epithelial colonization, and to determine whether non-CF and CF-derived epithelial tissues elicit individual responses upon *P. aeruginosa* infections, we performed bacterial infection assays using airway ALI models (Fig S1). These recapitulate the human lung epithelium, and are state-of-the-art within human airway infection models (*28*, *31*, *33*). Specifically, we evaluated infection capabilities and epithelial response, in ALI models generated with: 1) primary basal cells from non-CF (n = 3) or pwCF (n = 7) donors, and 2) the hTERT-immortalized basal airway cell line BCi-NS1.1 (*39*) (Fig S1). For each infection, we quantified epithelial response and bacterial proliferation by measuring: 1) the transepithelial electrical resistance (TEER) to quantify the integrity of the epithelial barrier, 2) the release of lactate dehydrogenase (LDH) to quantify the induced epithelial cell lysis, 3) the secretion of the chemokine Interleukin-8 (IL-8) to quantify the onset of inflammation, and 4) the bacterial colony-forming units (CFU) to quantify the bacterial load post-infection. Infections were carried out using a collection of early (*_E), intermediate (*_I), and late (*_L) *P. aeruginosa* clinical isolates from pwCF belonging to distinct clone types (lineages which differ by >10,000 SNPs), which comprise different stages of CF adaptation, evolutionary histories and infection scenarios (Table S1) (*40*, *41*). These clone types were chosen based on their frequency in the Copenhagen CF Clinic (DK01 and DK02) and worldwide (C and PA14 clone, DK06 and DK19 clones in Denmark, respectively). In addition, we included well-characterized *P. aeruginosa* reference strains (PAO1 and PA14), including a PAO1 derivative lacking the virulence determinant *pscC* of the Type III secretion system (T3SS). The T3SS plays an important role in the transition from acute to chronic/persistent infection occurring during host adaptive evolution (*42*, *43*). Additionally, it is involved in host colonization, invasion and cell damage, as we, and others have previously reported (*17*, *33*, *34*). The chosen bacterial strains exhibit distinct phenotypes characterized by both high and low growth rates, motility, and mild-to-high increased minimum inhibitory concentration (MIC) to tobramycin, ciprofloxacin, and ceftazidime (Fig S2). These strains are representative of the heterogeneity of *P. aeruginosa* infecting population present in pwCF (*44–46*). To ensure a comparable bacterial load post-infection across all samples, fast growing strains were incubated in ALI models for 14 h while slow growing strains were incubated for 38 h (Fig S3). Consistent with this experimental set-up, no significant differences in CFUs were observed across bacterial strains and ALI models (Kruskal-Wallis followed by Dunn’s post-hoc test, p>0.05) (Fig S3). Additionally, Pearson correlation analysis showed no correlation between the total CFU relative to TEER (*r*=-0.27, p=0.001), LDH release (*r*=-0.004, p=0.9601), or IL-8 secretion (*r*=0.12, p=0.1441), indicating independence of the variables. In addition, strain-specific bacterial infection outcomes (TEER, CFU, LDH, and IL-8) were overall comparable across non-CF, CF, and BCi-NS1.1 ALI models (Kruskal-Wallis followed by Dunn’s post-hoc test, p>0.05), except for DK01_L, DK06_L and DK19_L infections, where LDH release was significantly higher in CF compared to non-CF and BCi-NS1.1 ALI models (Kruskal-Wallis followed by Dunn’s post-hoc test, p<0.01, Table S2: Infection Dynamics). In all cases, Pearson correlation analysis of infection data across the different ALI models showed values > 0.9 (p<0.001) (Fig 1A). This suggests that in our model system, the infection outcomes (TEER, CFU, LDH, and IL-8) are mostly dependent on the specific virulence potential of the infecting bacteria rather than on the different ALI models (non-CF, CF, or BCi-NS1.1). All infection data are detailed in Table S2: Infection dynamics.

**Fig 1.**
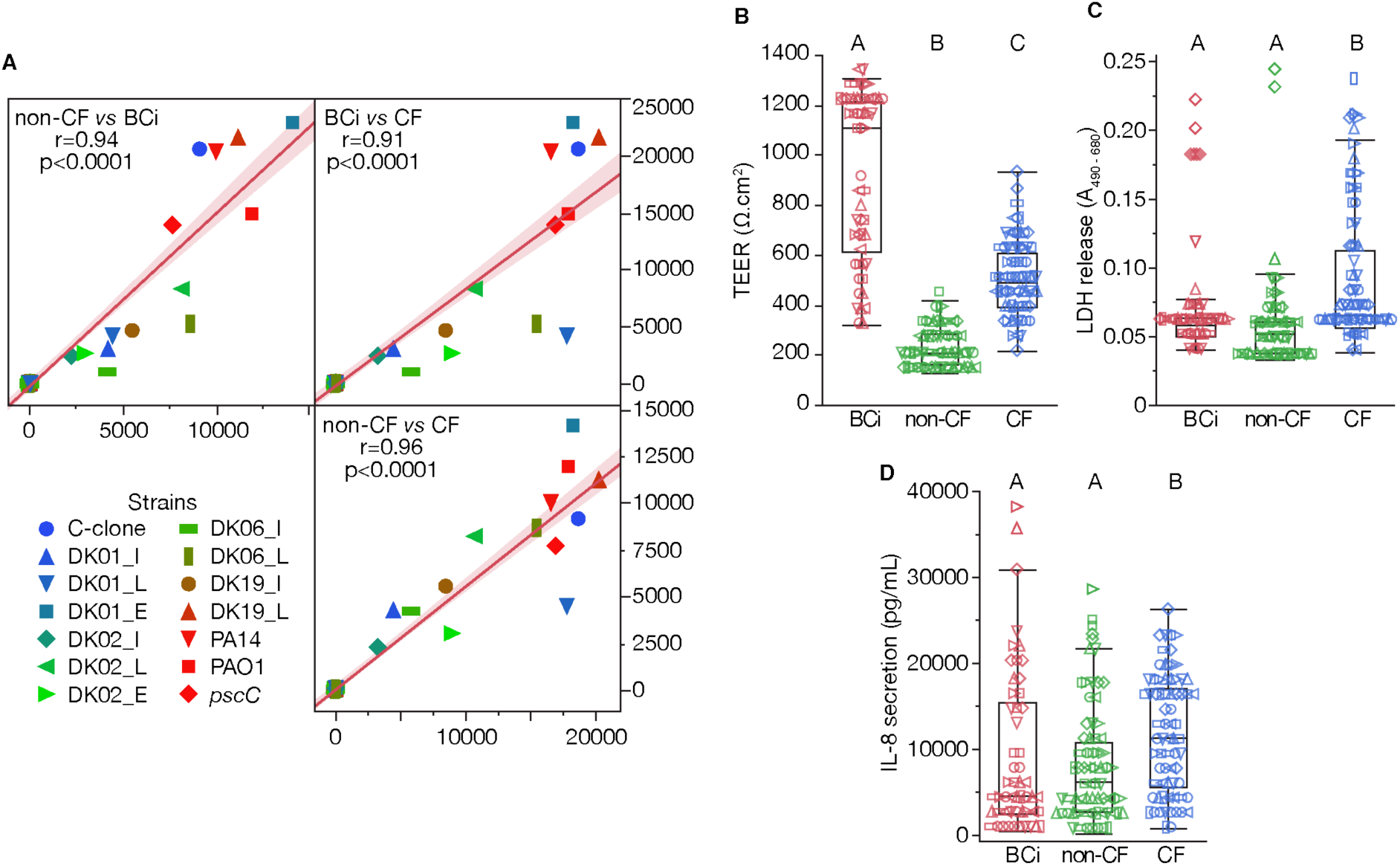
Infection outcomes across ALI models and bacterial strains. A) Pearson correlation analysis of the infection data across ALI models and bacterial strains. Pearson’s correlation coefficient r, statistical significance (p value), linear fitting between the variables, and 95 % confidence interval are indicated in the graph. B) TEER measurements of non-CF, CF, and BCi-NS1.1 ALI models after 28 days of differentiation in the absence of bacterial infections. Quantification of C) LDH release and D) IL-8 secretion by non-CF, CF, and BCi-NS1.1 models 14 h or 38 h post infection. Infections were carried out for 14 h for early and intermediate strains (DK01, DK02, DK06 and DK19) and laboratory strains (PAO1 and PA14) and for 38 h for late strains (DK01, DK02, DK06 and DK19). In B, C and D, statistical significance was computed by Kruskal-Wallis followed by Dunn’s post-hoc test, where connecting letters report significance between groups (p<0.05). CF: cystic fibrosis; BCi: BCi-NS1.1; TEER: transepithelial electrical resistance; LDH: lactate dehydrogenase; IL-8: Interleukin-8.

When comparing across all infections (ALI, n = 194), TEER measurements pre-infection revealed tissue dependent values, with BCi-NS1.1 models exhibiting the highest TEER followed by CF and non-CF ALI models (Kruskal-Wallis followed by Dunn’s post-hoc test, p<0.0001) (Fig 1B). Similarly, CF models exhibited the highest level of cellular damage (LDH release) and highest inflammatory response (IL-8 secretion) (Kruskal-Wallis followed by Dunn’s post-hoc test, p<0.01) (Fig 1C, D). Comparable results were obtained when discriminating by the duration of infection (14 h vs 38 h) and mock infected ALI models (Fig S4). Moreover, CF ALI models exhibited a time-dependent increase of LDH release (Kruskal-Wallis followed by Dunn’s post-hoc test, p<0.01) (Fig S4D).

The recent development of CFTR modulator treatment has changed the clinical approach to CF treatment. However, several reports indicate the persistence of *P. aeruginosa* within the airway of ETI treated pwCF (*10*, *12*), suggesting that restoration of airway epithelial functionality is not enough for pathogen eradication. For this reason, we tested *P. aeruginosa* infection capacity in ETI-treated and untreated CF ALI models. First, we confirmed the effect of ETI in CF ALI models (n = 3, donors homozygous for ΔF508) in the absence of *P. aeruginosa*. A significant decrease in TEER and an increase in transepithelial electrical conductance (TEEC), an indirect parameter of CFTR function (*47–49*), was observed in CF ALI models after 24 h of ETI treatment (Fig 2A, B). Contrary, no effect was observed in non-CF cells irrespective of ETI treatment (Fig S5A, B). Increased CFTR protein expression at the apical membrane was confirmed by immunofluorescent staining of ETI-treated CF ALI models (Fig 2C). Having confirmed restoration of CFTR function and localization, we next evaluated whether ETI treatment altered the infection capacity of *P. aeruginosa* in non-CF and CF ALI models (Table S2: ETI treatment). Infection outcomes in ETI-treated and untreated ALI models using PAO1 (Fig 2D, E) and the clinical strains DK01_I and DK02_L (Fig S5C, D) showed no difference in TEER, CFU, or bacterial distribution within the CF ALI models. A similar response was observed when the same strains infected non-CF ALI models (Fig S5E, F).

**Fig 2.**
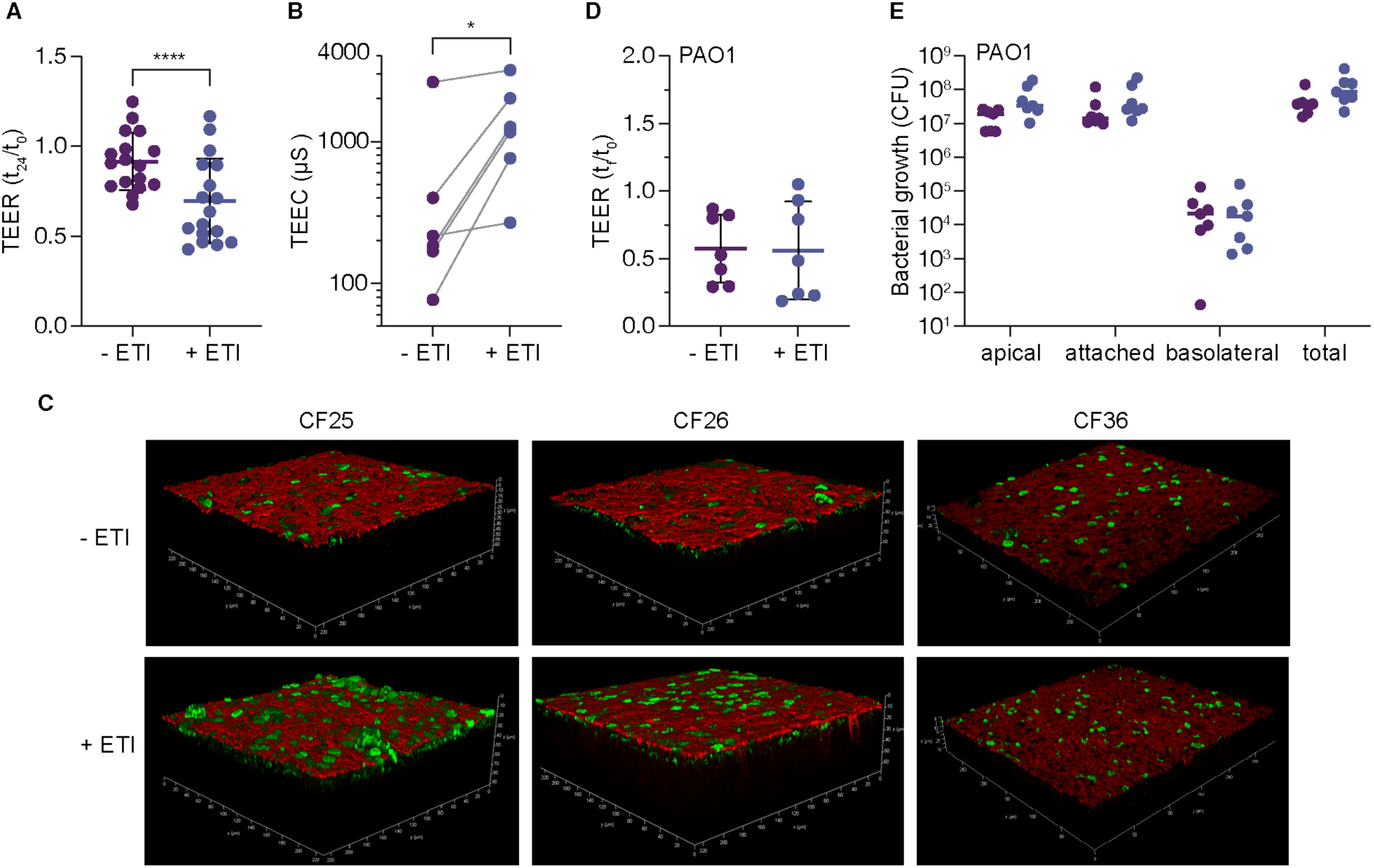
Effect of elexacaftor/tezacaftor/ivacaftor (ETI) on cystic fibrosis (CF) ALI models infected with PAO1. A) TEER fold change in CF ALI models treated ± ETI for 24 h. B) TEEC after CFTR stimulation in ALI models treated ± ETI. Each data point represents one independent donor. C) Three-dimensional reconstruction of confocal microscopy images of immunofluorescent staining of CFTR protein in ALI models from the CF donors ± ETI. Green: Anti-CFTR 570 (CFTR protein), Red: Phalloidin (F-actin). D) TEER change after 14 h of infection with PAO1 laboratory strain ± ETI. E) Bacterial growth (CFU) after 14 h of infection in the apical (non-attached / free bacteria), attached to the epithelium, and basolateral compartment, along with the total number of bacteria. In A, C and D, each data point represents one independent experiment, and the mean ± SD is included. Statistical significance was calculated by paired two-sided t-test where **p<0.01 and ****p<0.0001. ALI: air-liquid interface; TEER: transepithelial electrical resistance; TEEC: transepithelial electrical conductance; CFU: colony-forming units.

Despite intrinsic differences between non-CF, CF, and BCi-NS1.1 ALI models, our results suggest that infection dynamics are independent of the ALI type or modulator treatment and may depend on the bacterial lineage or other factors not recreated in the ALI model.

### Strain-specific infection phenotypes of *P. aeruginosa*

After years of adaptive evolution in pwCF, *P. aeruginosa* changes its phenotype in response to stresses such as antibiotics, reactive oxygen species (ROS), immune cells and others (*24*, *50*). Therefore, we next evaluated whether differences in strain adaptive evolution and historical contingency affected the infection capabilities in a convergent manner. Given that bacterial infections progressed similarly across the different ALI models and independent of ETI treatment, for each strain, we averaged the infection data from non-CF, CF, and BCi-NS1.1 ALI models (Table S2: Infection Dynamics Median.).

PCA and K-Means clustering identified four clusters of strains with similar infection properties which can be associated primarily with their virulence, epithelial damage, and localization within the epithelium (Fig 3A). PC 1 (44.8 % of the variance) clusters strains based on their virulence potential with cluster 1 and 2 showing a decrease in TEER, and an increase in cytotoxicity (LDH release), and inflammatory response (IL-8) (Fig 3A, B). Contrary, PC 2 (36.2 % of the variance) separates strains based on their ability to penetrate the epithelium. Strains from cluster 1 are found in the basolateral compartment, indicating increased penetration through the epithelium, while strains from cluster 3 are found predominantly attached within the epithelium (Fig 3A, B). It is worth noting that while the *pscC* mutant strain, and the DK02_L isolate, group in the more virulent cluster 1 and 2, respectively, both strains show intermediate infection capabilities comparable with strains from cluster 3 and 4 (Fig 3B). Interestingly, while virulent strains from cluster 1 and 2 colonized the epithelia as large aggregates/biofilm structures, less virulent strains from cluster 3 and 4 infected as single colonies or small bacterial aggregates, as shown by confocal microscopy (Fig 3C and Fig S6). Furthermore, most virulent strains (PAO1, PA14, and DK02_L) and a few low virulent strains (*pscC* mutant strain and DK06_L isolate) caused cell shedding. Lastly, variations in colonization pattern were observed within each cluster. For instance, from cluster 2, only PA14 was found within the epithelium, whereas DK01_E was found overlapping with mucus-producing cells. The latter was also seen for DK01_I and DK02_I from cluster 4 (Fig 3C, Fig S6).

**Fig 3.**
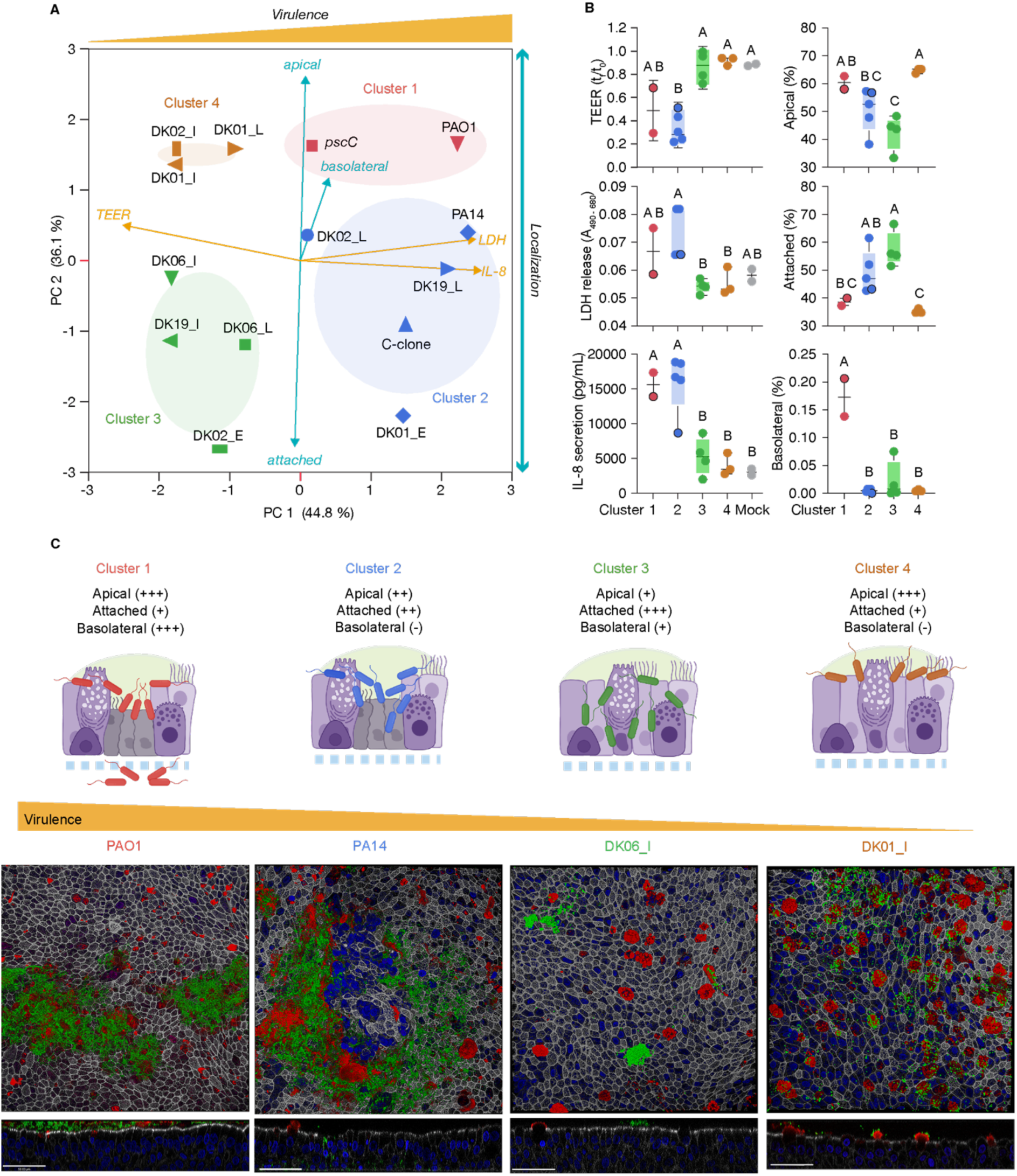
Infection phenotypes of *Pseudomonas aeruginosa* strains. A) PCA and K-mean clustering of the infection phenotypes of laboratory (PAO1, *pscC* mutant strain, PA14, and C-clone) and clinical strains (early, intermediate, and late DK01, DK02, DK06, and DK19) of *P. aeruginosa.* Each symbol represents the median infection phenotypes in non-CF, CF, and BCi-NS1.1 ALI models explained by the variables TEER, LDH release, IL-8 secretion, and distribution of bacteria in the apical, attached, and basolateral compartment. K-mean clusters (1 to 4) and PCA loadings are color-coded to underline groupings of strains and explained variance, respectively. B) Box plot of the specific K-mean clusters characteristics for TEER, LDH release, IL-8 secretion, and normalized distribution of CFUs in the apical, attached, and basolateral compartment. The outlined symbols in clusters 1 and 2 represent *pscC* mutant strain and DK02_L strains, respectively. Statistical significance was computed by Kruskal-Wallis followed by Dunn’s post-hoc test, connecting letters report significance between groups (p<0.05). C) Colonization phenotype of strain-specific clusters in CF cells. Schematic representation of CF ALI models (purple) challenged with strains from clusters 1 (red), 2 (blue), 3 (green), and 4 (brown). Representative confocal microscopy images of CF ALI models infected with strains from Cluster 1 (PAO1), Cluster 2 (PA14), Cluster 3 (DK06_I) and Cluster 4 (DK01_I). Top and cross-sectional views of ALI models are shown. Representative images from all strains infecting CF ALI models are shown in Fig S6. Immunofluorescent staining: sGFP (green) staining chromosomally tagged bacteria; MUC5AC (red) specific for mucus producing cells; Phalloidin (white) specific for actin, and DAPI (blue) nuclear stain. Stainings were done in three biological replicates, one representative image is shown for each infection. PCA: principal component analysis; TEER: transepithelial electrical resistance; LDH: lactate dehydrogenase; IL-8: Interleukin 8.

### Dual RNA-sequencing of host and pathogen in *P. aeruginosa*-infected ALI models

While different bacterial strains showed similar infection patterns across all ALI models, the molecular response of both bacteria and host is likely influenced by the specific comparisons (*i.e.*, non-CF, CF, or BCi-NS1.1 ALI models and different *P. aeruginosa* strains). To test this hypothesis, we performed dual RNA-seq analysis of ALI models from the BCi-NS1.1 cell line, a non-CF donor, and a CF, with PAO1, *pscC* mutant strain and mock infected (Fig 4A, Table S2: RNAseq). This approach allowed us to analyze the molecular responses of isogenic bacteria, while capturing differences in the colonization process of highly virulent and intermediate virulent-like clinical strains, represented by the PAO1 and *pscC* mutant strain, respectively (Fig 3A). While it is known that the T3SS is crucial in host colonization and invasion (*33*), its role at the molecular level during host interactions remains unknown.

**Fig 4.**
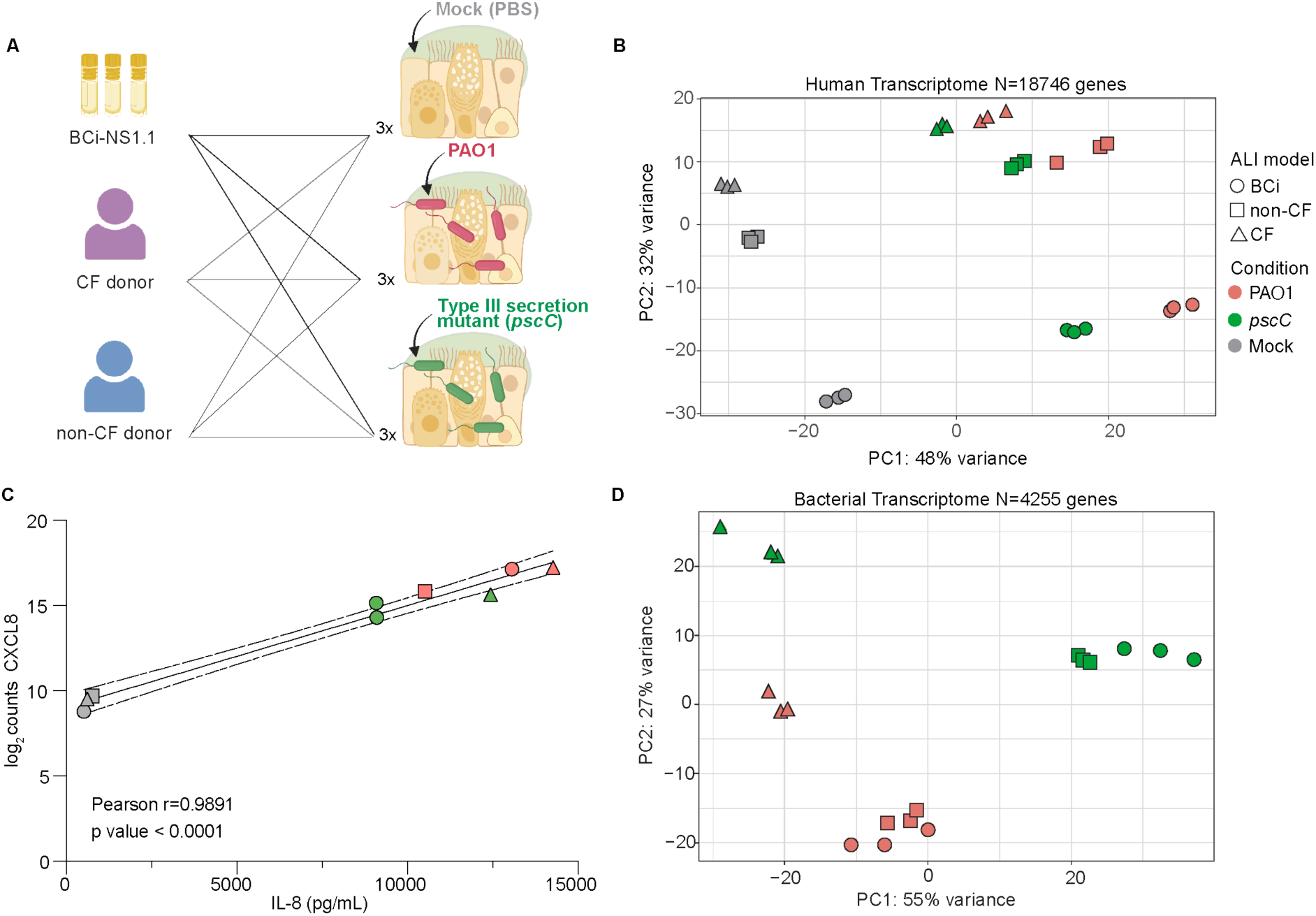
Exploring the transcriptional responses from human and pathogen upon infection. A) Schematic representation of the ALI models (non-CF, blue; CF, purple; BCi-NS1.1, yellow) and bacteria (PAO1, red; *pscC* mutant strain, green) used to perform infections and subsequent dual-RNAseq. PCA of log_2_ read counts from B) the host and D) the pathogen. Data include the reads of 18746 genes from the human transcriptome and 4255 from the *Pseudomonas aeruginosa* transcriptome, from which differential gene expression analysis was performed. The type of ALI model is presented by symbol shapes: BCi-NS1.1 (circle), CF (triangle), and non-CF (square). Mock infected samples are colored in grey, whereas ALI models infected with PAO1 or *pscC* mutant strain are shown in red and green, respectively. Each data point represents one biological replicate. C) Regression lines and Pearson’s correlation coefficient for IL-8 quantification and CXCL8 gene expression (log_2_ read counts). 95 % confidence interval lines for the linear regression fit are included. ALI: air-liquid interface; CF: cystic fibrosis; PCA: Principal component analysis; TEER: transepithelial electrical resistance; IL-8: Interleukin-8.

PCA analysis of the host response (n = 18746 genes) showed that: 1) the transcriptional program of infected ALI models (PAO1 and *pscC* mutant strain) differed from mock infected (mock) (Fig 4B), and 2) that primary cells, *i.e.*, non-CF and CF ALI models, exhibited a different host response relative to BCi-NS1.1 models (Fig 4B). Pearson correlation analysis of gene expression (n = 18746) in mock infected non-CF, CF, and BCi-NS1.1 ALI models, was positive and significant (Pearson’s r>0.97; p<2.2e^-16^), thus ensuring that any intrinsic variability between the different models would not significantly impact the downstream differential expression analysis (Fig S7A). A similar trend was found in mock infected samples, when only analysing the 455 differentially expressed genes (DEGs) found in the comparison PAO1 vs mock (Pearson’s r>0.94; p<2.2e^-16^) (Fig S7B), indicating that the different ALI models do not have functional impact on gene regulation post infection. IL-8 secretion and CXCL8 gene expression, responsible for IL-8 protein expression, was positively correlated (Pearson’s r = 0.9891; p<0.0001) confirming higher levels of chemokine secretion in infected samples compared to mock infected samples, thus validating our RNA-seq data (Fig 4C).

Analysis of bacterial expression profiles (n = 4255 genes) revealed: 1) bacteria infecting non-CF and BCi-NS1.1 ALI models behave similarly, 2) infection in CF models drives a distinct bacterial expression profile, and 3) differences dependent on the bacterial genetic background (Fig 4D).

### Cellular and inflammatory response in ALI models upon infection with PAO1 and *pscC* mutant strain

To characterize the molecular basis of the host response to either PAO1 or *pscC* mutant strain, we performed individual DEG comparisons between ALI models infected with A) PAO1 and mock (Fig 5A), and B) PAO1 and *pscC* mutant strain (Fig 5D). For each comparison, we first compared gene convergence/divergence across the different cell types and second, we performed gene set enrichment analysis using both KEGG and the MSigDB (GO) database.

**Fig 5.**
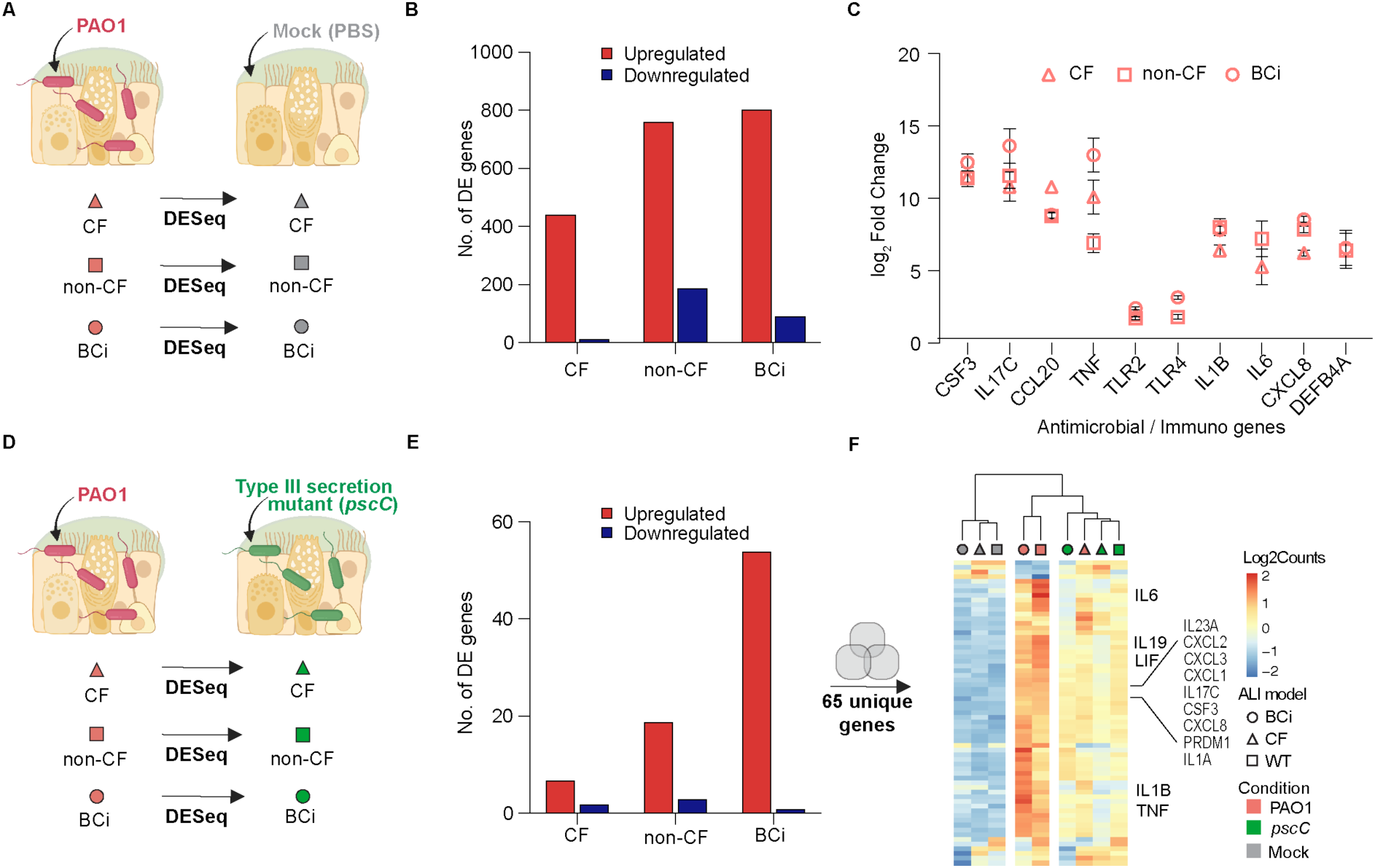
Intrinsic host response before and after colonization with PAO1 and *pscC*. A) Representation of the individual differential expression comparisons between ALI models were performed: PAO1 and mock infected ALI models. B) Number of differentially expressed genes (DEGs) shown as upregulated (red) and downregulated (blue). An adj p value < 0.05 and log_2_ Fold Change ≥ |1| was used as cutoff C) log_2_ Fold Change of immune genes when comparing PAO1 infected and mock. Only significantly expressed DEGs are shown. D) Representation of the DEG comparisons between PAO1-infected and *pscC* mutant strain-infected ALI models. E) Number of DEGs shown as upregulated (red) and downregulated (blue). F) Heatmap of log_2_ read counts scaled for each gene (row) from the unique 65 DEGs from D) Each column represents the mean of the three biological replicates sequenced clustered based on the result of pvclust analysis. CF: cystic fibrosis; BCi: BCi-NS1.1.

Compared to mock, CF ALI models showed the fewest DEGs, followed by non-CF and BCi-NS1.1 ALI models, when infected with PAO1. For all non-CF, CF, and BCi-NS1.1 infections, we observed mainly upregulated genes (Fig 5B, Table S3). Enrichment analysis showed that *P. aeruginosa* triggered upregulation of several pathways related to inflammation via nuclear factor-kappa B (NF-κβ) and tumor necrosis factor alpha (TNF*α*) and overexpression of cytokines, including IL-6 and IL-17 (Table S4). Of note, colony-stimulating factor (CSF3) and pro-inflammatory cytokine IL17C were among the most DEGs across all types of ALI models (mean log_2_ FoldChange = 12.01 ± 1.5 and 11.8 ± 0.6, respectively) (Fig 5C). Other highly expressed genes included chemokine ligand 20 (CCL20) and TNF*α*, Toll-like Receptor (TLR)-2, TLR4 (involved in recognition of bacterial pili and LPS), IL1B, IL6, CXCL8 (involved in the first stages of infection) and the antimicrobial human beta-defensin 4 (DEF4BA) (Fig 5C).

We found subtle differences in host response following infection with the less virulent *pscC* mutant strain, compared to PAO1 (Fig 5E). In fact, only 65 unique genes were differentially expressed in ALI models infected with PAO1 relative to *pscC* mutant strain (Fig 5E). Pathways related to cytokine signaling and inflammation were overrepresented among this set of genes, which were primarily highly expressed in PAO1-infected non-CF and BCi-NS1.1 ALI models (Fig 5F). This suggests that, at the time tested, the delivery of effectors from the T3SS partially affects host gene expression. However, *P. aeruginosa* has a wide array of other virulence factors contributing to infection and concomitant inflammatory response, possibly explaining the subtle difference observed between the PAO1 and *pscC* mutant strain.

### PAO1 and *pscC* mutant strains elicit differential response upon interaction with ALI models

Analysis of the bacterial transcriptome aimed to evaluate how the expression profile of PAO1 and *pscC* mutant strain vary upon interaction with different ALI models (Fig 6A). As previously shown (Fig 4C), whole-gene expression profiling of both bacterial strains was similar in non-CF and BCi-NS1.1 ALI models. Indeed, no PAO1 gene was differentially expressed when comparing non-CF vs BCi-NS1.1 infections, and a similar observation was made for *pscC* mutant strain with only three DEGs (Fig 6B top and bottom panel, Table S5). The PAO1 transcriptome (n = 4255 genes) showed only 56 and 79 DEGs in CF vs non-CF and CF vs BCi-NS1.1 infections, respectively, suggesting that PAO1 behaves similarly when colonizing different ALI models (Fig 6B). Contrary, *pscC* mutant strain showed a 3.9-fold and 5-fold increase in DEGs, in CF vs non-CF, and CF vs BCi-NS1.1 infections, respectively (Fig 6B, bottom panel). This suggests that the activity of T3SS has a direct impact on the bacterial expression profile upon infection in CF ALI models.

**Figure 6.**
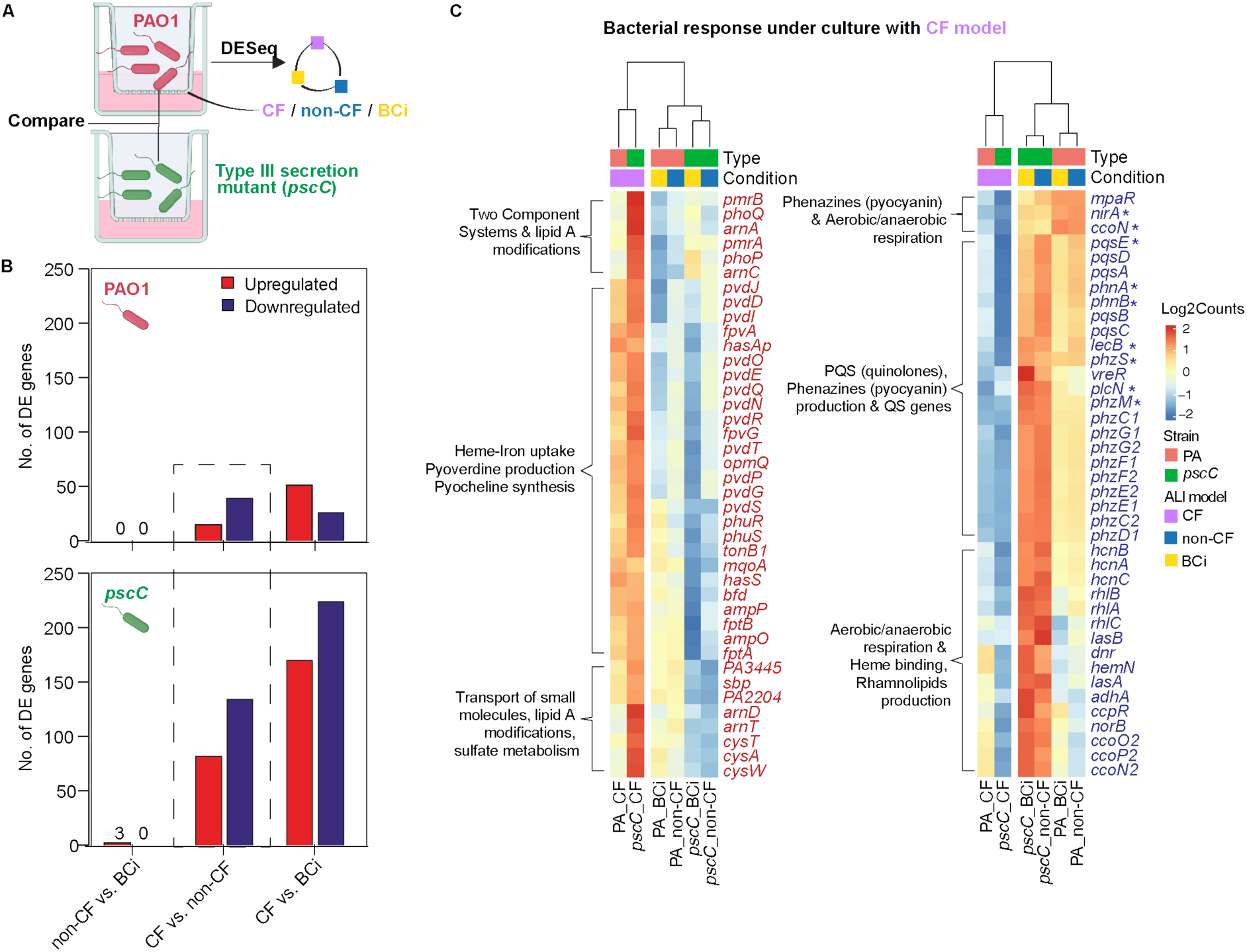
Bacterial response in different ALI models. A) Representation of the differentially expressed gene (DEG) comparison of PAO1 and *pscC* mutant strain transcriptomes analyzed in different ALI models, *i.e.*, non-CF vs. BCi-NS1.1, CF vs. non-CF, and CF. vs BCi-NS1.1. B) Number of DEGs shown as upregulated (red) or downregulated (blue) and separated by strain (PAO1 top and *pscC* mutant strain bottom panel, respectively). An adj p value < 0.05 and log_2_ Fold Change ≥ |1.0| was used as cutoff. C) Heatmap of log_2_ read counts scaled for each gene (row) selected from categories from Table S6, comparison *pscC* CF vs. non-CF. Each column represents the mean of the three biological replicates sequenced, clustered based on the result of pvclust analysis. Genes with asterisk (*) were differentially expressed in both PAO1 and *pscC* mutant strain, when comparing CF vs. non-CF models. CF: cystic fibrosis; BCi: BCi-NS1.1.

Considering similarities between bacterial gene expression in non-CF and BCi-NS1.1 ALI models, we next compared the expression profiles of *pscC* mutant strain and PAO1, focusing on CF and non-CF infections (Fig 6B underlined boxes). Compared with PAO1, an increased number of genes was regulated in the *pscC* mutant strain upon infection in CF ALI models (PAO1 n = 55, *pscC* mutant strain = 218) (Fig 6B). A total of 41 genes were common and included upregulation of genes involved in respiration under iron-limiting conditions such as 1) *bo*_3_ quinol oxidase (*cyo*) genes and 2) members of the *fumC-sodM* iron-responsive operon (Table S6). Likewise, downregulation of several pathways involved in virulence, such as the production and transport of phenazines (among others: *phz1* and *phz2* operons), nitrate reductase *nirA,* and fucose-specific lectin *lecB* was seen in both strains (*53–55*).

When comparing the infection of *pscC* mutant strain in CF vs non-CF ALI infections, we found enrichment of three main categories of upregulated genes: 1) iron chelation (related to heme uptake, pyoverdine, and pyochelin transport), 2) sulfate metabolism (*cysAWT-sbp* operon) and, 3) lipid A modifications (Fig 6C, left panel and Table S6). Contrary, several genes related to virulence, including QS and biofilm pathways, were enriched in the downregulated class (Fig 6C, right panel, and Table S6). For example, 1) genes from the *pseudomonas* quinolone signal (PQS) cluster, 2) rhamnolipids coding genes, 3) the type 6 secretion system H2 gene cluster (H2-T6SS), 4) genes from oxide-sensing mechanisms and, 5) elastases (*lasB*) and pyocyanin biosynthetic genes. This demonstrates that the bacterial response differs depending on the host environment.

In summary, our results show that *P. aeruginosa* regulates several pathways involved in iron starvation, respiration, and virulence mechanisms in a CF ALI model, in response to different environmental cues. Many of these traits have been previously linked to adaptation *in vivo* CF airways (*56*, *57*), thus underscoring the robustness of our infection model.

## Discussion

Persistent bacterial infections are often associated with antibiotic resistance. However, several other mechanisms, such as reduced virulence, slow growth, and immune evasion, have been proposed to explain antibiotic treatment failure (*3*). This underscores the need to understand the infection and colonization processes at the host-pathogen interface beyond antibiotic resistance. Several *in vitro* studies of clinical strains have shown convergence toward low virulent phenotype as chronic infection progresses (*22*, *25*, *40*, *41*). However, molecular changes often poorly reflect *in vivo* conditions due to differing environment and absence of host interactions (*56*, *57*). Recently, specific mutations in *P. aeruginosa* were discovered to be involved in novel adaptive mechanisms that transcend conventional antibiotic resistance, highlighting the need for host-focused approaches to develop more effective strategies for treating bacterial infections (*34*, *38*).

To address this, we quantified infection outcomes during the early stages of colonization using different clinical isolates of *P. aeruginosa* by comparing three types of ALI infection models (non-CF, CF, and BCi-NS1.1). We quantified host response including epithelial integrity (TEER), cellular cytotoxicity (LDH), inflammatory response (Interleukin 8), bacterial load (CFU), and localization within the ALI models (confocal microscopy) to provide systematic insight into the infection capabilities of *P. aeruginosa*. Irrespective of the type of ALI model (non-CF, CF, and BCi-NS1.1), colonization was predominantly driven by the intrinsic bacterial virulence potential. Infection clusters were neither based on the genetic background of the strains (*i.e.*, isolates from the same lineage were not always grouped together), nor on the level of adaptation (*i.e.*, late strains did not always show reduced virulence). While high virulence has been identified as a key driver during the early stages of host colonization (clusters 1 and 2), the preferential bacterial localization, either attached to the epithelium (cluster 3) or planktonic (apical) growth (cluster 4), has not previously been shown in clinical isolates. This distinction is highly relevant, since bacterial localization has significant implications for treatment failure (*38*). In our ALI model using CF-derived primary cells, we observed similar infection profiles, regardless of ETI modulator treatment or bacterial strain. This is in agreement with recent works showing that, although there is a reduction of bacterial load in pwCF treated with ETI, bacteria persist in the airway (*10*, *12*). The observed trends provide valuable ground knowledge about the inherent properties of the ALI models employed, as well as about the virulence of the infecting bacteria. Importantly, the ALI model holds predictive potential as it allows evaluation of drug efficacy, immune system activity, host response, and inter-species interactions, which are fundamental to understanding infection progression and improving treatment outcomes.

Recent transcriptomics studies have explored *P. aeruginosa* interactions in ALI models (*30*, *32*, *36*, *58*). However, comparative insight into the molecular responses of non-CF, CF, and BCi-NS1.1 ALI models remains limited. To address this gap, we performed dual transcriptomic analysis of *P. aeruginosa* infections across these model systems. Given the biological complexity and technical challenges associated with primary cells, our analysis focused on infections with the reference strain PAO1 and a less virulent *pscC* mutant, since the T3SS plays a key role in early host colonization and invasion (*33*).

Importantly, a high correlation between the three types of ALI models was observed and only 2.4 % of the identified genes were differentially expressed across all comparisons. This indicates that the primary response and gene expression profiles are comparable between ALI models independently of the host origin or donor. Similar to earlier research, we found that colonization of host epithelium with *P. aeruginosa* triggers activation of inflammatory pathways, response to bacteria and LPS through upregulation of NF-kβ and TNF*α* signaling pathways and overexpression of cytokines including IL-6 and IL-17 signaling (*30*, *36*). In agreement with the latter, infections carried out with eight different donors (n = 3, non-CF and n = 5, CF) showed similar trends in IL-8 secretion for PAO1 across the different ALI models. On the other hand, deletion of the *pscC* gene induced fewer host DEGs related to inflammation compared to PAO1. This aligns well with infection readouts, where the *pscC* mutant strain exhibited intermediate virulence between PAO1 and the more adapted clinical strains.

Upon infection and depending on the availability of nutrients and/or host signals, *P. aeruginosa* tightly controls the regulation of a wide array of virulence factors involved in host-mediated respiratory infections (*59*). In our CF ALI model, both PAO1 and *pscC* mutant strain limited the expression of *lasB*, a principal extracellular virulence factor, and the biosynthesis of pyocyanin, both contributing to inflammation and epithelial disruption (*60*). The downregulation of these genes may enable the bacteria to evade the immune system and reduce host inflammation, as observed in our infected CF ALI models, which showed fewer DEGs (n = 442) compared to non-CF (n = 761) and BCi-NS1.1 ALI models (n = 804) models. Strikingly, the interaction between CF ALI models and the *pscC* mutant strain showed a more complex regulation resembling many of the features already seen in clinical CF isolates. Besides phenazine biosynthesis, the *pscC* mutant strain also showed downregulation of the PQS system and rhamnolipid production, mirroring key features of CF late chronic strains (*56*). Another example is the high expression of genes involved in the production and transportation of siderophores pyoverdine (*pvd*) and pyochelin (*pch*), as well as the assimilation of heme (*phu* and *has* systems) (*57*), suggesting limited accessibility of these micronutrients in CF ALI models relative to non-CF.

While infection models allow for a more representative analysis of bacterial phenotypes, they also have some limitations. It was, indeed, surprising that only a few isolates showed significant differences in one or two infection outcomes between non-CF and CF ALI models. This result may be influenced by several factors: a) the ALI model does not fully replicate the *in vivo* lung environment, which is influenced by the presence of immune cells, potential antibiotic exposure, and oxygen and micronutrient gradients, b) the chosen infection time points (14 or 38 h) may be sub-optimal for detecting subtle host response patterns, c) the enclosed nature of the ALI transwells may limit mucociliary clearance, or d) the absence of a residing airway microbiota, which could modulate host-pathogen interactions. Therefore, further investigations are required to systematically evaluate the contribution of each of these parameters.

Altogether, we explored the virulence potential of clinically persistent clones of *P. aeruginosa* when colonizing different airway epithelial tissues grown at the ALI interface. We covered a variety of bacterial lineages and host donors and defined clusters of infection properties. Furthermore, we quantified the global transcriptional response of both host and pathogen upon infection. Our data foster the need for studies using host-pathogen models to recapitulate the *in vivo* conditions of pathogenic diseases. Moreover, given the more appropriate physiological environment, ALI models serve as a platform for evaluating treatment efficacy and identifying novel therapies for the treatment of persistent infections. Yet, further studies implementing immune cells, antibiotic treatment, and other infecting species are needed to understand infection progression, antibiotic treatment failure, and new mechanisms of persistence.

## Materials and Methods

### Ethical approval and consent to participate

This study was approved by the local ethics committee of the Capital Region of Denmark (Region Hovedstaden), registration number H-20024750. All patients provided informed consent to participate in the study and for the publication of associated results and pertinent clinical data. Pseudonymization of patient information was performed to untie any links that could lead to patient identification.

### Primary basal cell extraction, cell lines and culture

Primary basal cells were obtained from nasal brushes, nasal polyps, and distal lung biopsies from pwCF (n = 10) and non-CF (n = 5) donors. Details on sample source and CFTR mutation are provided in Table S2 separated according to assay.

Nasal brushes were obtained with a 2.5 mm cytology brush and nasal polyps, and lung biopsies were surgically removed. All samples were placed in fresh culture media, supplemented with primocin (Invivogen, ant-pm-05), gentamicin/amphotericin solution (Gibco, Cat #10184583) and pen/strep (Gibco, Cat #15140122), immediately following collection. Nasal brush samples were homogenized by resuspension, whereas nasal polyps and lung biopsies were disrupted by scraping and dissection to release basal cells. To remove any blood and mucus, nasal polyps and lung biopsies were washed three times in fresh media. Next, samples were centrifuged at 300 x g for 5 min at room temperature (R/T) and cell pellets were resuspended in PneumaCult™-Ex Plus Medium, with PneumaCult™-Ex Plus 50X (STEMCELL Technologies, Cat #05040), according to manufacturer’s instructions, supplemented with 96 ng/mL hydrocortisone (STEMCELL Technologies, Cat #07925) and 10 µM ROCK inhibitor (Y-27632) (Tocris Bioscience Cat #1254), hereafter ExPlus media. BCi-NS1.1, Basal Cells Immortalized-NonSmoker 1.1, an hTERT expressing airway cell line, and primary cells (non-CF and CF), were seeded into culture flasks (Greiner, item no. 690160 & item no. 658175) and maintained in ExPlus media. When reaching 80 % confluence, cells were passaged by detachment with TrypLE (Gibco Cat #12563011) for further expansion or for seeding into transwells for ALI culture. Specifically for primary cells (non-CF and CF), cell culture flasks were pre-coated with human type-I collagen (Gibco, Cat #A1048301), ExPlus media was further supplemented with antibiotics as described above, and only passage 1-3 were used for ALI models. All cells were maintained at standard conditions, 37 °C, 5 % CO2 in a humidified incubator.

### Air-liquid interface (ALI)

To establish ALI models, 1.5 x 10^5^ basal cells from non-CF, CF, and BCi-NS1.1 were seeded in transwells (6.5 cm^2^, 0.4 µm pores) (Corning Life Sciences product no. 3470) pre-coated with human type-I collagen (Gibco, Cat #A1048301). ALI models were cultured in ExPlus media, prepared as described, added to the apical and basolateral compartment. Upon full confluence, ALI was established by aspirating the media from both compartments, and adding PneumaCult™-ALI Medium, with PneumaCult™-ALI 10X Supplement and PneumaCult™-ALI Maintenance Supplement (100X), according to manufacturer’s instructions (STEMCELL Technologies, Cat #05001), supplemented with 480 ng/mL hydrocortisone (STEMCELL Technologies, Cat #07925) and 4 µg/mL heparin (STEMCELL Technologies, Cat #07980) to the basolateral compartment. ALI models were cultured for a minimum of 28 days, media was refreshed as required. When necessary, ALI models were rinsed apically with PBS to remove excess mucus. ALI models were maintained at standard conditions, 37 °C, 5 % CO2 in a humidified incubator.

### ETI modulator treatment of infected and noninfected ALI models

On day 28 of ALI culturing, a combined CFTR-modulator treatment of 3 μM elexacaftor (VX-445, MedChemExpress), 10 μM tezacaftor (VX-661, Selleck Chemicals LLC), and 5 μM ivacaftor (VX-770, Selleck Chemicals LLC), hereafter ETI, was added for 24 h to the basolateral medium. ETI compounds were prepared fresh before each experiment. Untreated controls were treated with an equal volume of DSMO. For infection studies, ALI models were treated ± ETI for 24 h prior to infection, as described above. Infections were carried out with PAO1 (14 h), DK01_I (14 h), and DK02_L (38 h), adding fresh ETI at the start of infection.

### Transepithelial electrical conductance (TEEC) measurement

TEEC is calculated from the reciprocal value of TEER measurements:

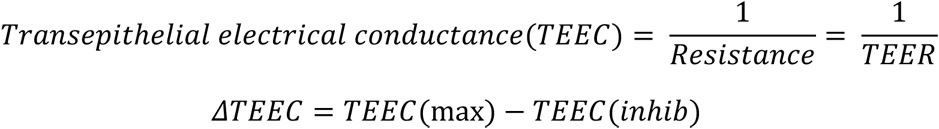

TEER was measured after maximal CFTR stimulation with forskolin (TargetMol, Cat #T2939) and genistein (TargetMol, Cat #T1737), and subsequent inhibition with PPQ-102 (TargetMol, Cat #T1874), before converted to TEEC. First, Coon’s modified Ham’s F12 medium without NaHCO_3_ (DiagnoCine, Cat #AT146), supplemented 20 mM Na-Hepes (SigmaAldrich) (pH 7.3), was added to the apical and the basolateral compartment, and equilibrated for 1 h, after which, a baseline TEER was measured. Following, 10 µM amiloride (SigmaAldrich, Cat #A7410) was added to the apical compartment. Next, apical and basolateral medium was refreshed, and 10 μM forskolin and 50 μM genistein was added. Subsequently, a solution containing 30 μM PPQ-102 (CFTR inhibitor) was added to the apical compartment. Between each step, a 10 min incubation time was allowed, after which TEER was measured, and subsequently converted to TEEC. ALI models, non-CF and CF, were tested in at least three technical replicates.

### Clinical strains, growth conditions and fluorescent tagging

Strains used in this study are summarized in Table S1. Clinical strains were obtained from sputum samples from pwCF attending or who have attended the Copenhagen Cystic Fibrosis Center at University Hospital Rigshospitalet, Copenhagen, Denmark. Sputum sampling is part of the routine visits to the CF clinic and was not performed for the purpose of this research. All clinical strains except DK01_E and DK02_E, were previously sequenced and their genomic and transcriptomic profiles characterized (*40*).

*P. aeruginosa* reference strains PAO1, PA14, and C-clone were obtained from the Leibniz Institute DSMZ collection with references DSM22644, DSM19882, and DSM29238, respectively. Additionally, an isogenic *pscC* mutant strain (PAO1 background), previously constructed and used for infection studies in our lab was included (*17*).

All strains were tagged with a superior green fluorescent protein (sfGFP) to enable visualization by confocal microscopy. Bacterial strains were GFP-tagged by 4 parental mating using the miniTn7 delivery plasmid with rhaSR-PrhaBAD inducible promoter (pJM220) (*61*). The plasmid was engineered to accommodate the sfGFP gene. Transformants were selected on *Pseudomonas* isolation agar supplemented with 30 µg/mL gentamicin (SigmaAldrich, Cat #G3632). The final sfGFP-tagged strains were verified by fluorescence using a Leica DM4000 B epifluorescence microscope and growth rate (Fig S8).

### Phenotypic characterization of *P. aeruginosa* strains

GFP-tagged strains were characterized based on their growth in synthetic cystic fibrosis media (SCFM), prepared as previously described(*62*), bacterial movement, and antibiotic susceptibility profiles. Briefly, single colonies were picked and grown on liquid Luria-Bertani broth (LB) cultures. Cultures were standardized to OD_600_ = 0.1 and further diluted to OD_600_ = 0.01 in SCFM media and growth was recorded every 15 min for 18 h in a 96 well plate reader at 37 °C with continuous shaking. Doubling times and growth rate estimations were calculated using Graph Pad Prism software (v 10.2.2), focusing on the exponential part of the growth curve. For swimming assay, 2 µL of the OD_600_ = 0.1 cultures were plated on the center of tryptone broth plates (10 g/L tryptone, 5 g/L NaCl) with 0.25 % agarose. Motility diameter was measured after incubation for 6 h at 37 °C and additional 16 h, at R/T. For antimicrobial susceptibility, the microdilution method from EUCAST was used (The European Committee on Antimicrobial Susceptibility Testing 2023). Cultures were standardized to Mc Farland (5 x 10^5^ CFU/mL) and grown in Müller-Hinton broth with serial fold dilutions of either tobramycin (SigmaAldrich, Cat #T4014), ceftazidime (SigmaAldrich, Cat #C3809), or ciprofloxacin (SigmaAldrich, Cat #17850) for 16 h in 96 well plates at 37 °C with continuous shaking. MIC was calculated as the concentration of antimicrobial required for complete growth inhibition of the strain as detected by the naked eye. All phenotypic experiments were performed with three independent cultures.

### Infection of ALI models with *P. aeruginosa*

Overnight *P. aeruginosa* cultures were standardized to OD_600_ = 0.05 (reference, early, and intermediate strains) or OD_600_ = 0.1 (late strains) and grown until mid-exponential phase in 5 mL of LB medium. Subsequently, 1 mL of culture was centrifuged at 5000 × g for 5 min, and bacterial pellets were washed and resuspended in PBS at a concentration of 1 x 10^7^ CFU/mL. For each strain, the OD:CFU ratio was tested in triplicates to achieve reproducible results. A final of 1 x 10^3^ CFU was used to inoculate the apical side of ALI models. Mock infected samples were inoculated with an equal volume PBS. Infected ALI models were incubated at 37 °C, 5 % CO_2_ in a humidified incubator for the specified duration required to characterize the infection (14 h for reference, early, and intermediate strains, and 38 h for late strains). At the end of infection, 200 µL of PBS was added apically, and TEER was measured.

Subsequently, apical PBS and basolateral media was collected for CFU quantification by plating 10 µL of serial dilutions on LB-agar plates. Remaining sample was preserved at −80 °C for downstream analysis of LDH and IL-8. For quantification of attached bacteria, 200 μL of Triton X-100 (0.1 % in PBS), with no harm to bacteria (*34*), was added apically, the epithelium disrupted using a cell scraper, and CFU were quantified as described above.

### Transepithelial electrical resistance (TEER)

Transepithelial electrical resistance (TEER) was measured prior to and immediately at the end of infection using the EVOM2 (World Precision Instruments) and the STX2 electrode (World Precision Instruments). The electrode was submerged into 200 µL PBS at the apical compartment and 400 µL media in the basolateral compartment.

### Immunofluorescence staining and confocal microscopy

ALI models were fixed in 4 % paraformaldehyde for 20 min, at 4 °C. Following, ALI models were washed 3 times with PBS, and incubated in blocking buffer (0.1 % Triton X-100, 1 % saponin, 3 % BSA in PBS) for 1 h, at R/T. Next, ALI models were incubated with primary antibodies diluted in staining solution containing 3 % BSA and 1 % saponin in PBS at 4 °C, overnight. Next day, ALI models were washed 3 times in PBS and incubated with secondary antibodies and dyes, diluted in staining solution, for 2 h, at R/T. After staining, membranes were excised from the transwells and mounted onto microscope slides with VECTASHIELD® Antifade Mounting Medium (VWR, Cat #VECTH-1000). Images were acquired with the Leica Stellaris 8 Confocal Microscope (40X, Oil objective) and analyzed using the LasX software 1.4.4.26810 (Leica) and Imaris 10.1.0 software.

### Antibodies and dyes

The following antibodies and dyes were used: anti-MUC5AC (Invitrogen Cat #MA5-12178, 1:100), anti-CFTR 570 (kind gift from cystic fibrosis foundation, 1:100), Phalloidin-Alexa Fluor 555 (Invitrogen Cat #A30106, 1:400). Alexa Fluor-647 Goat anti-mouse (Invitrogen Cat #A21235, 1:500) and DAPI nuclear dye (Thermo Scientific Cat #62248, 1:750).

### Lactate dehydrogenase (LDH) assay

LDH release was determined with the CyQUANT™ LDH Cytotoxicity Assay Kit (Invitrogen, Cat #C20301) according to manufacturer’s instructions. For this, 50 µL of basolateral media was taken from ALI models, and absorbance was measured at 680 nm and 490 nm. All samples were analyzed in triplicates.

### Interleukin-8 (IL-8) quantification

IL-8 secretion was measured with the Human IL-8/CXCL8 DuoSet ELISA kit (Bio-Techne, Cat #DY208) according to manufacturer’s instructions. Basolateral media was diluted 1:10 and absorbance was measured at 450 nm and 540 nm. All samples were analyzed in triplicates.

### RNA extraction from ALI cultures and sequencing library preparation

Basal cells derived from a lung biopsy of a non-CF donor (non-CF 16), a nasal brush from one pwCF (CF14), and the BCi-NS1.1 cell line were used to generate ALI models. These models were infected with either PAO1, *pscC* mutant strain, or mock infected in six independent transwells. ALI models and infections followed the same procedures as described for other samples in this study. Infection outcomes were consistent with the experimental variability observed across donors (Table S2).

Total RNA was extracted using the Qiagen RNeasy Plus Mini kit, following the manufactureŕs protocol. Briefly, after infection and TEER measurements, PBS was removed from the apical compartment, and 300 µL of RNA protect Cell Reagent was added. Adherent cells were detached by pipetting without the need of enzymatic dissociation and two transwells from the same condition were pooled per 300 µL of sample and frozen at −80 until RNA extraction. For enzymatic lysis, samples were thawed, centrifuged to remove RNA protect, then resuspended in 1 mL of RNA-se free water containing 1 mg/mL of lysozyme, and incubated for 30 min at 37°C. Samples were then centrifuged and resuspended in 600 µL RLT Buffer (with β-mercaptoethanol), and RNA was extracted following the Qiagen protocol to recover fragments >200 nt. RNA was quantified using the Qubit® RNA BR Assay kit (Invitrogen), and integrity assessed with an Agilent 2100 Bioanalyzer using the RNA Nano kit. All samples had DV_200_ values (percentage of RNA fragments >200 nt) ranging from 50 % to 75 % and were included in downstream processing.

2 µg of eluted RNA were treated with DNAse I (Invitrogen) for 30 min at 37°C in a final volume of 50 µL. The treated RNA was purified using Qiagen columns by combining 150 µL RLT Buffer (without β-mercaptoethanol), 150 µL 95-100 % ethanol, and 50 µL RNA sample. Following two washes with RPE buffer, RNA was eluted in 20-25 µL RNA-se free water.

A total of 500 ng of RNA per sample was used to prepare sequencing libraries using the Illumina Stranded Total RNA Prep with Ribo-Zero Plus kit. Sequencing was performed on an Illumina NextSeq 500 system using a high output kit, generating a minimum of 30 million (M) single end 75 bp reads per sample.

As bacterial transcripts represent only a small fraction of the total RNA within host cells (*63*), RNA was extracted from two independent infected transwells and combined to form a single biological replicate. Each condition included three such replicates, and no more than 12 samples were pooled for sequencing run. This approach enhanced the recovery of bacterial RNA, yielding between 2.5 M to 10 M reads per sample depending on the bacterial strain and cell type (Fig S9, Table S7). For samples infected with the *pscC* mutant strain (non-CF-*pscC* 1-3, CF-*pscC* 2-3, BCi-*pscC* 1-3), a second sequencing round was performed to achieve a minimum threshold of 1 M reads mapping to the *Pseudomonas* genome. Table S7 lists the sequencing yields for both runs (BCi-*pscC* 1a and BCi-*pscC* 1b, etc.).

### Host-pathogen transcriptome analysis

Sequencing reads were analyzed with our in-house pipeline updated for host-pathogen samples (*64*). Briefly, low-quality bases and contaminant adapters were trimmed using Trimmomatic (v 0.35), discarding reads shorter than 35 nt. Reads were further processed using the SortMeRNA tool (v 2.1) to remove reads generated from residual rRNA transcripts. High-quality human and bacterial reads were separated *in silico* by mapping reads using the BWA aligner and MEM algorithm against the human genome assembly GRCh38.p9 retrieved from the NCBI database. Reads not mapping on the human genome were used as input for analyzing the transcriptionally active *P. aeruginosa* population mapping against *P. aeruginosa* PAO1 genome (NCBI: NC_002516.2) using BWA (0.7.15-r1140) aligner and MEM algorithm with default parameters.

### Gene expression analysis

Mapped reads from both host and *P. aeruginosa* were quantified using htseq-count (v 0.7.2). Resulting count tables were imported into R for differential gene expression (DEG) analysis. For host data, genes with ≥10 reads in at least three samples were retained, resulting in a filtered dataset of 18746 genes. Counts were normalized across all samples using the variance stabilizing transformation (VST-normalization) in the DESEq2 package (v 1.44.0) with the option blind = TRUE (*65*). For bacterial reads, only genes with ≥10 reads in all replicate samples were retained, yielding 4255 *P. aeruginosa* genes. VST-normalized counts were used for exploratory analysis including principal component analysis (PCA) using `plotPCA()`function, visualized with `ggplot()`, and hierarchical clustering analysis (HCA). When necessary, correlation plots were generated from normalized count data using Pearson’s correlation coefficients, with significance values reported. Heatmaps of selected gene sets were generated using `pheatmap()`, based on condition-wise replicate means.

DEGs were identified from DESeq2 results using the `results()` function with the `contrast` argument for pairwise comparisons. Genes were considered differentially expressed with a log₂ Fold Change ≥ |1.0|and an adjusted p value ≤ 0.05.

Significant host DEGs were analyzed for gene set enrichment analyses with the “Human Hallmark” collection from the Molecular Signature Database (MsigDB). Gene sets were retrieved with `msigdbr()`, and enrichment was performed using the `GSEA()` with the following parameters: `minGSSize=5`, `maxGSSize=500`, `pvalueCutoff=0.05`, `pAdjustMethod=”BH”`. Additional enrichment analysis using the KEGG database was performed using `gseKEGG()` under the same parameters. When overlapping pathways were identified (e.g., “response to lipid” vs. “cellular response to lipid”), the term with the lowest adjusted p value was selected.

For bacterial DEGs, functional enrichment was conducted using log_2_ Fold Change values and gene-term associations from COGs and KEGG annotations based on *P. aeruginosa* PAO1 data from Pseudomonas.com, via `GSEA()` and `gseKEGG()` with `organism=”pae”`.

To assess overlap among DEG lists, UpSet plots were generated using the UpSet() function in R.

## Data sharing

Raw sequence data supporting the results of this work are available as SRA BioProject under Accession No. PRJNA1153754 and accession numbers from individual BioSamples and sequences is summarized in Table S7. All data is available in the publication.

## Author Contribution Statement

**Claudia Antonella Colque:** Methodology, Formal analysis, Investigation, Data curation, Visualization, Writing – original draft, Writing – review & editing. **Signe Lolle:** Methodology, Formal analysis, Investigation, Writing – review & editing. **Ruggero La Rosa:** Formal analysis, Data curation, Visualization, Writing – original draft, Writing – review & editing. **Maria Pals Bendixen:** Investigation, Writing – review & editing. **Asha M. Rudjord-Levann:** Formal analysis, Writing – review and editing. **Filipa Bica Simões:** Methodology, Formal analysis, Investigation, Writing – original draft, Writing – review & editing. **Kasper Aanæs**: Resources. **Søren Molin:** Conceptualization, Supervision, Writing – review & editing. **Helle Krogh Johansen:** Conceptualization, Funding acquisition, Resources, Supervision, Writing – review & editing.

## Acknowledgments

The authors thank the Danish pwCF and the non-CF donors who participated in this study, Laura Elisabeth Bojesen Haapalo and Rene Horsleben Petersen (Department of Cardiothoracic Surgery, Copenhagen University Hospital, Rigshospitalet, Denmark) for providing and assisting with lung tissue samples, and the nurses from Rigshospitalet CF clinic assisting with patient recruitment and sample collection. We thank the cystic fibrosis foundation (CFF) for providing the CFTR anti570 antibody, and Ronald G. Cristal (Weil Cornell Medical College, New York, USA) for providing the BCi-NS1.1 cell line. This research was funded by a Challenge Grant (Ref. nr.: NNF19OC0056411) and a grant from The John and Birthe Meyer Foundation awarded to HKJ. HKJ and SM were supported by a grant from CAG - Greater Copenhagen Health Science Partners (GCHSP), 2020. The work at the Novo Nordisk Foundation Center for Biosustainability (CfB) is supported by the Novo Nordisk Foundation (Ref. nr.: NNF10CC1016517). We thank the Infection Microbiology group (Oihane Irazoqui Sanchez, Pablo Laborda Martínez, Janus Anders Juul Haagensen) for their insightful comments and discussions. We thank Elio Rossi (Department of Biosciences, University of Milan, Italy) for his advice on transcriptomics analysis, and Nicoletta Pedemonte (IRCCS Institute Gaslini, Genova, Italy) for her assistance in modulator treatment assays.

